# Occurrence of (oo)cysts of *Cryptosporidium* and *Giardia* on vegetables across Nepal

**DOI:** 10.64898/2026.04.06.716709

**Authors:** Retina Shrestha, Bhanu Bhakta Neupane, Basant Giri

**Author notes:** Correspondence: Basant Giri, P. O. Box: 23002, Kathmandu, Nepal Bhanu Bhakta Neupane.

## Abstract

Gastrointestinal disorder caused by the ingestion of (oo)cysts of *Cryptosporidium* and *Giardia* is one of the major health problems in developing countries. Fruits and vegetables that are usually consumed unpeeled, poorly washed and or cooked and are the major modes of transmission. Frequent large-scale screening of the food samples is necessary to prevent outbreaks but screening of vegetables for such microbes is limited in Nepal. In this study, we used a smartphone microscopy system to study prevalence and quantification of (oo)cysts of *Cryptosporidium* and *Giardia* in 651 vegetable samples collected from nine major vegetable collection sites across Nepal. The overall prevalence rate of vegetable samples was 37.5% with at least with one of the parasites. We found that 23.2% samples were contaminated with *Giardia* and 33.3% samples were contaminated with *Cryptosporidium*. Among eight vegetable types, the prevalence rate was lowest in carrot (20%) and highest in spinach (48%). The prevalence rate of vegetable samples at different sites ranged from 13% in Dhading to 61% in Dhangadi. The contamination rate was 28% for winter, 43% for summer and 33% for monsoon seasons in samples collected from Kathmandu. These vegetables should be considered as a potential source of parasitic contamination in people. These vegetables can cause infection if consumed poorly washed and or cooked, posing a potential source of parasitic contamination in people.

## 1. Introduction

Food-borne illnesses have been rising globally in past several decades. According to the World Health Organization (WHO)’s fact sheet on food safety, globally 1 in 10 people fall ill after consuming contaminated food resulting in up to four hundred thousand deaths each year. Children under 5 years of age are more vulnerable as they carry 40% of the foodborne disease burden (WHO, 2022). The impact of these illnesses is skewed towards low- and middle-income countries (LMICs) including Nepal with an annual estimated cost of USD 110 billion productivity accounting the losses and medical treatment costs(WHO, 2022). The skewness may be because of poor hygiene, poor health care and reporting systems, and a lack of affordable diagnostic facilities and infrastructures.

*Cryptosporidium parvum* and *Giardia duodenalis* are two of the widespread protozoan parasites responsible for intestinal infections. They cause cryptosporidiosis and giardiasis resulting in chronic diarrhea and digestive disorders. Both protozoans are obligate and intracellular parasites that attach to epithelial cells lining of the digestive and respiratory tracts of a wide range of hosts (Cacciò, Thompson, McLauchlin, & Smith, 2005). Infection of these protozoans hinders the barrier function and absorption capability leading to mild-to-severe diarrhea, weight loss, stunting, malnutrition, abdominal cramps, and cognitive impairment (Donowitz et al., 2016; Korpe et al., 2016). Usually, infants, young children, immunocompromised individuals, farmers, animal handlers, and international travellers are at peril (Swaminathan et al., 2009; Swirski, Pearl, Peregrine, & Pintar, 2016). It is also very common to find people with asymptomatic infection of *Cryptosporidium* and *Giardia* in the endemic areas of developing countries. Furthermore, these protozoans have a low infectious dose (10-1000 for *Cryptosporidium parvum* and 10-100 for *Giardia duodenalis*) (Smith, Caccio, Cook, Nichols, & Tait, 2007) and have a high excretion rate, ranging from >5 ×10^3^ to 9.2× 10^5^ oocysts per gram faeces for *Cryptosporidium* and 5.8× 10^5^ cysts per gram faeces for *Giardia* (Danciger & Lopez, 1975; Goodgame, Genta, White, & Chappell, 1993). The infective stage of *Cryptosporidium* oocysts is spherical with 4-6 µm in diameter and *Giardia* cysts are oval with 8–12 µm long and 7–10 µm wide. This microscopic size and low specific gravity of infective stages facilitate their dissemination in food and water. Additionally, the (oo)cysts are robust and resistant in the environment. Therefore, they can remain infective for weeks on stored produce, and have zoonotic and anthroponotic potential (Abeywardena, Jex, & Gasser, 2015).

Infections with *C. parvum* and *G. duodenalis* are acquired by the ingestion of their oocyst and cyst, respectively from contaminated food and water. The (oo)cysts contaminate the food on the surface. Fruits and vegetables that are usually consumed unpeeled, raw or partially cooked, and improperly washed are a major risk factor and one of the major modes of transmission. The complex and porous surface of vegetables and fruits helps pathogens to attach on the surface and survive. The contamination of food products can occur directly by (oo)cysts in faeces from humans and animals, through the soil and water, cross-contamination from other vegetables during storage and transportation, and by food handlers (Al-Binali, Bello, El-Shewy, & Abdulla, 2006; Amoah, Drechsel, Abaidoo, & Klutse, 2007; Pires, Vieira, Perez, Wong, & Hald, 2012; Tefera, Biruksew, Mekonnen, & Eshetu, 2014a). The production, transportation and supply chain of vehicles require cool and moist conditions to maintain the freshness of the food products which facilitates the survival of (oo)cysts. Additionally, simple washing with distilled water does not completely remove *Cryptosporidium* oocysts from the surface of the vegetables as they are highly resistant to washing and are very well attached to the peel. To completely remove the parasites, vegetables should be washed with chemicals like calcium hypochlorite (Cook & Lim, 2012; Fallah, Pirali-Kheirabadi, Shirvani, & Saei-Dehkordi, 2012).

Nepal is one of the least developed countries with a population of approximately 29 million. A significant Nepali population (∼33.8%) reside in rural municipalities and 4.5% households do not have toilet (National Statistics Office, Government of Nepal, 2023). Consumption of contaminated drinking water is common which directly augment the chances of getting diarrheal diseases. According to the Nepal Demographic and Health Survey 2022 (Ministry of Health and Population [Nepal], New ERA, and ICF, 2023), the occurrence of diarrhoea in children under five years old was 10%. In the year 2017 alone, 1193 death of five-year-old children by diarrhoea was reported in Nepal, which was higher than in other South Asian countries like Bangladesh. Intestinal parasitic infections are common in Nepal and ranks fourth in ‘top-ten-diseases’ in the country. The lower socio-economic status and uneducated population are among the most vulnerable groups in the country(S. K. Rai, Hirai, Abe, & Ohno, 2002).

Several studies have shown the prevalence of these protozoans in human samples in Nepal. In a study carried out at Kanti Children’s Hospital, Kathmandu in 2002, *Giardia lamblia* (12%) was the most common parasite studied among children aged 1-15 years (K. Rai, Sherchand, & Bhatta, 2004a). 10.4% children of the age group 0-15 years were infected with *C. parvum*.

A 7.4% prevalence of *G. lambia* among children of age group 0-15 years was reported in Lalitpur district. Around 50% of the diarrheal diseases among children in Kathmandu and rural hills were found to be due to intestinal parasites and prevalence ranged from 32.6% to 72.4% among them (K. Rai, Sherchand, & Bhatta, 2004b; S. K. Rai et al., 2004). Infection with *Giardia duodenalis* has been reported from other locations such as Dolakha, Solukhumbu, Dadeldhura, Pokhara, Birgunj. In addition to human samples, few reports have shown the prevalence of these protozoans in sewage water, river water, and cattle(Mahato, Singh, Rana, & Acharya, 2018; Sherchand, Ghimire, & Mishra, 2005).

It is important to identify the source of protozoans’ contamination for better prevention of diseases. However, there are limited data on the occurrence of *Giardia* and *Cryptosporidium* contamination in food and drinking water sources in Nepal. One of the limiting factors for large scale monitoring of food and water samples for microbial contamination is the unavailability of affordable analytical tools. In our previous publication, we addressed the problem by developing a low-cost smartphone microscopic system that can detect both *Giardia* and *Cryptosporidium* (oo)cysts in vegetable and water samples (Shrestha et al., 2020). In this paper, we report on the results of testing 651 vegetable samples for (oo)cysts contamination using the smartphone microscope. The eight types of vegetables that are mostly consumed raw were collected during 2019 – 2020 from nine vegetable markets located in eight districts of Nepal and tested them.

## 2. Materials and Method

### 2.1. Sampling sites and sample collection

Nepal is an agricultural country. Agriculture contributes 23.9% in the country’s GDP (Government of Nepal, 2022). However, Nepal also imports food items from India, China, Vietnam, Thailand, Spain, Italy, Australia, Indonesia. In 2017 alone, Nepal imported 706 thousand tonnes of vegetables, mostly from India which accounted for nearly 40% of the total demand of the central vegetable market in Kalimati (CASA, 2020).

Nepal has three broad ecological regions running from north to south, classified as Mountain (>3000 m), Hill (1000-3000m), and Tarai (<1000m). Rapid changes in altitude creates a wide range of climatic conditions in Nepal experiencing four seasons - summer, monsoon, autumn, winter, and spring. Summer season is very humid and records a highest temperature up to 40 °C. It is followed by monsoon season which is characterised by high precipitation. About 80 percent of annual precipitation falls under this time. (Nepal Tourism Board, 2026) We collected 651 vegetable samples during 2019 – 2020 from nine vegetable markets located in in eight districts of Nepal as shown in figure 1. The sampling sites represented four out of seven provinces of Nepal: Bagmati province (Kathmandu, Dhading, and Chitwan), Lumbini province (Rupendehi and Banke), Karnali province (Salyan and Surkhet) and Sudarpaschim province (Attariya, Kailali). Name of the local vegetable markets are Kalimati Vegetable Market, Kathmandu; Balkhu Vegetable Market, Kathmandu; Agriculture Product Collection Center, Dhading; Mahanagar Fruits and Vegetable Market, Bharatpur, Chitwan; Agriculture Product Wholesale Bazar, Butwal, Rupandehi; Agriculture Product Market Center, Kapurkot, Salyan; Bulbule Chhetriya Krishi Upaj Bazar, Surkhet; Agriculture Product Wholesale Market Center, Kohalpur, Banke; and Attariya Agriculture Product Wholesale Market, Attariya, Kailali. We collected samples from Kalimati fruits and vegetable market, Kathmandu in three different seasons representing monsoon, winter, and summer. The samples were purchased from retailers and wholesalers by secrete shoppers without informing the shopkeeper the purpose of purchase. These markets sell domestic as well as imported vegetables, usually from India. Vegetables from these markets are regularly distributed to the local markets and other cities in Nepal as well.

**Figure 1:**
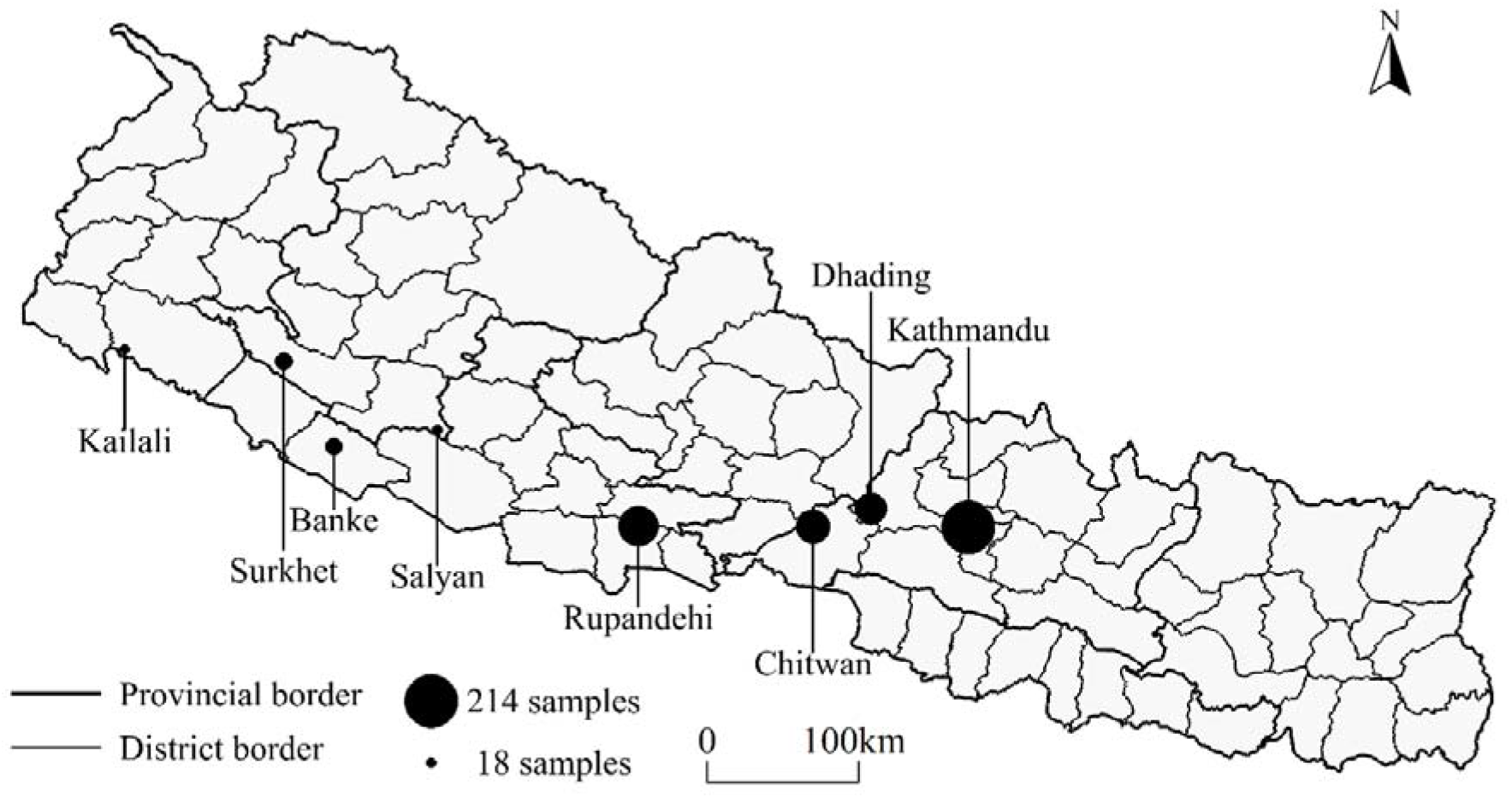
Map of Nepal showing sample collection sites. Size of the circles in each site indicate the number of samples collected

The vegetable samples included tomato (n=117), carrot (n=66), cabbage (n=98), radish (n=70), cucumber (n=97), green chili (n=112), spinach (n=41), and green onion (n=19).

Details of the samples is provided in Table S1. These vegetables are mostly consumed in raw form, have the potential for faecal contamination, have substantial contact with the ground, and need intensive fertilization and irrigation during cultivation. Around 0.5 to 1 kg of each sample were purchased from random shops in the vegetable markets. After the purchase, the samples were transferred separately in transparent plastic bags, labelled with sample ID and date of collection, and transported to the laboratory in an ice box.

### 2.2. Sample Processing

The vegetable samples were processed according to the Environmental Protection Agency method 1623 and *Codex Alimentarius* CAC/GL 33 with some modifications as described in our previous publication (Shrestha et al., 2020).

To ensure the method to be representative each sample was carefully mixed and reduced by quartering sampling technique. Three replicates of each vegetable were randomly selected to minimize sampling error and stored in a zip lock bag at 4 °C to 10 °C in refrigerator until extraction. The tray in which the vegetables were placed was wiped with 70% ethanol each time before starting new set of vegetable to avoid cross-contamination(Shrestha et al., 2020).

Once the vegetable samples were reduced by the quartering sampling technique, the weight of each vegetable sample were recorded. The average weight of vegetables such as tomato, cucumber, carrot, and radish were nearly 80-100g, cabbage and spinach were approximately 25g, while green chilies and green onion were around 15g.

Extraction of (oo)cysts from the surface of vegetables involved washing the sample with 1M glycine buffer at pH 5.5. The eluates were then carefully transferred into falcon tubes. The (oo)cysts in the eluate suspension were then concentrated by using the triple centrifugation method at 1500 x g for 10 minutes. Then the pellets were collected for microscopic analysis.

### 2.3. Microscopic examination of (oo)cysts

We used a handheld smartphone-based microscopic system developed in our laboratory to test the samples. The performance of the system is comparable with commercial brightfield and fluorescence microscopes as described in our earlier publication (Shrestha et al., 2020). In brief, the microscope used a 1mm diameter ball lens attached to the rare camera of the smartphone and commercial white LED for illumination. The microscope provided ∼200X magnification and ∼200 µm clear FOV. The sensitivity and specificity of the microscope were 67% and 93% for *Giardia* and 88% and 86% for *Cryptosporidium*, respectively (Shrestha et al., 2020).

For testing the vegetable extract, 10 µL of the concentrated extract was stained with 10 µL of Lugol’s iodine (1:2 in water) (HiMedia, Mumbai, India) and subsequently loaded in a hemocytometer (Max Levy, Philadelphia, USA). After incubating the samples for 6 minutes, the (oo)cysts were screened and enumerated in four quadrants of the hemocytometer. Triplicate measurement was made for each concentrated suspension. Average of triple measurement was used to estimate the number of (oo)cysts in the sample using recovery efficiency data previously estimated for different vegetables. The prevalence rate was estimated by the ratio of contaminated sample to total number of samples.

### 2.4. Statistical analysis

Statistical analysis and plots reported in this paper were performed using R studio version 4.0.3. Pair wise proportional test was used to compare the rate of contamination among different seasons, locations, and vegetables. Bonferroni post-hoc test was used to adjust p-value. The difference was considered significant at P<0.05. We chose this non-parametric analysis because the data did not meet the assumption of normality and data transformations failed to achieve normality.

## 3. Results and Discussion

### 3.1. Occurrence of *Giardia* and *Cryptosporidium*

Out of total vegetable samples, 37.5% (244) on average were contaminated with at least one (oo)cyst. The occurrence of *Giardia* and *Cryptosporidium* out of total contaminated samples is shown in figure 2 and Table S2. We found that 23.2% samples were contaminated with *Giardia* and 33.3% samples were contaminated with *Cryptosporidium*. The difference between the occurrence of singular cysts, singular oocysts, and both (oo)cysts was found to be significant (chi-square test: df=2 P<0.05).

**Figure 2:**
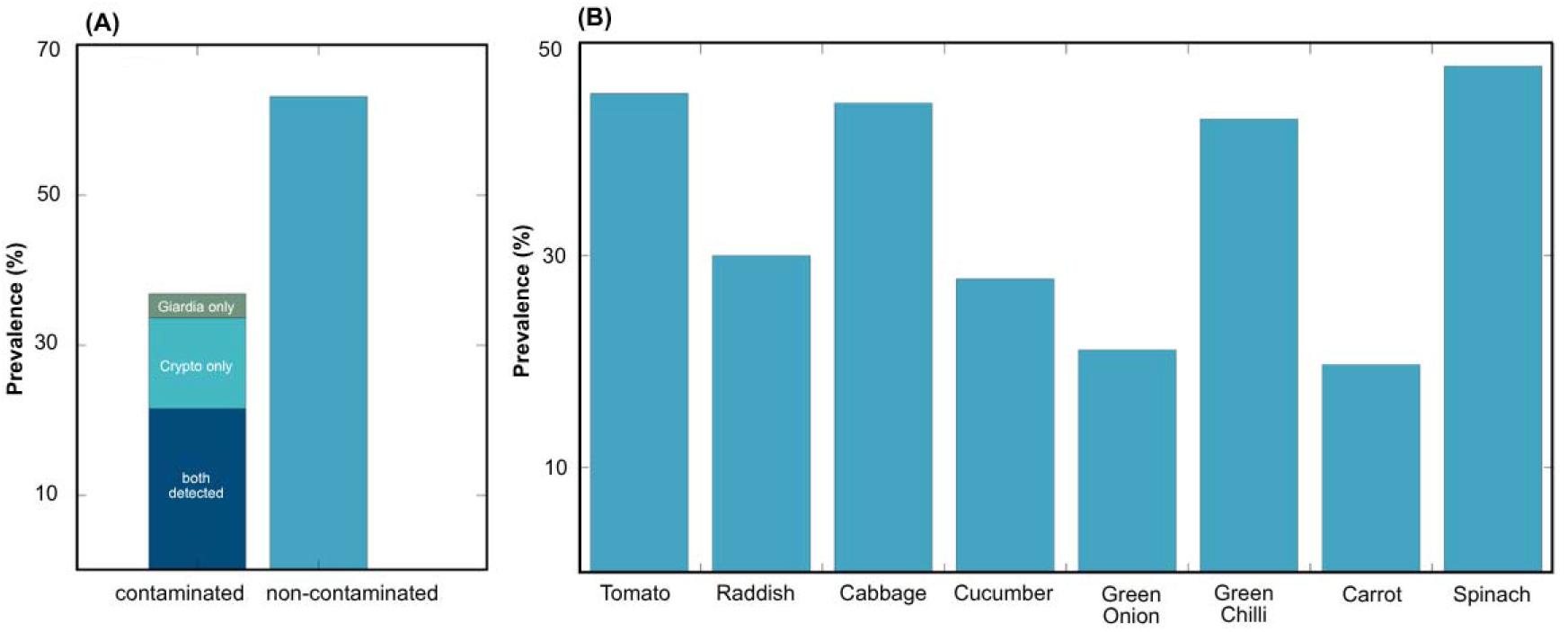
Prevalence of the (oo)cysts in aggregate samples (A) and in different types of vegetables tested (B).

The contamination of vegetables and fruits with *Cryptosporidium* oocysts and *Giardia* cysts has been documented in many countries (*see* table 1). The wide range of prevalence among different areas could be due to several factors which include geographic location, type of vegetable, number of samples, methods used for detection, type of water used for irrigation, quality of manure and post-harvesting handling method. In addition, population-related hygienic habits, sanitary facilities and climatic conditions also affect the contamination rate (Eraky, Rashed, Nasr, El-Hamshary, & Salah El-Ghannam, 2014b).

**Table 1:**
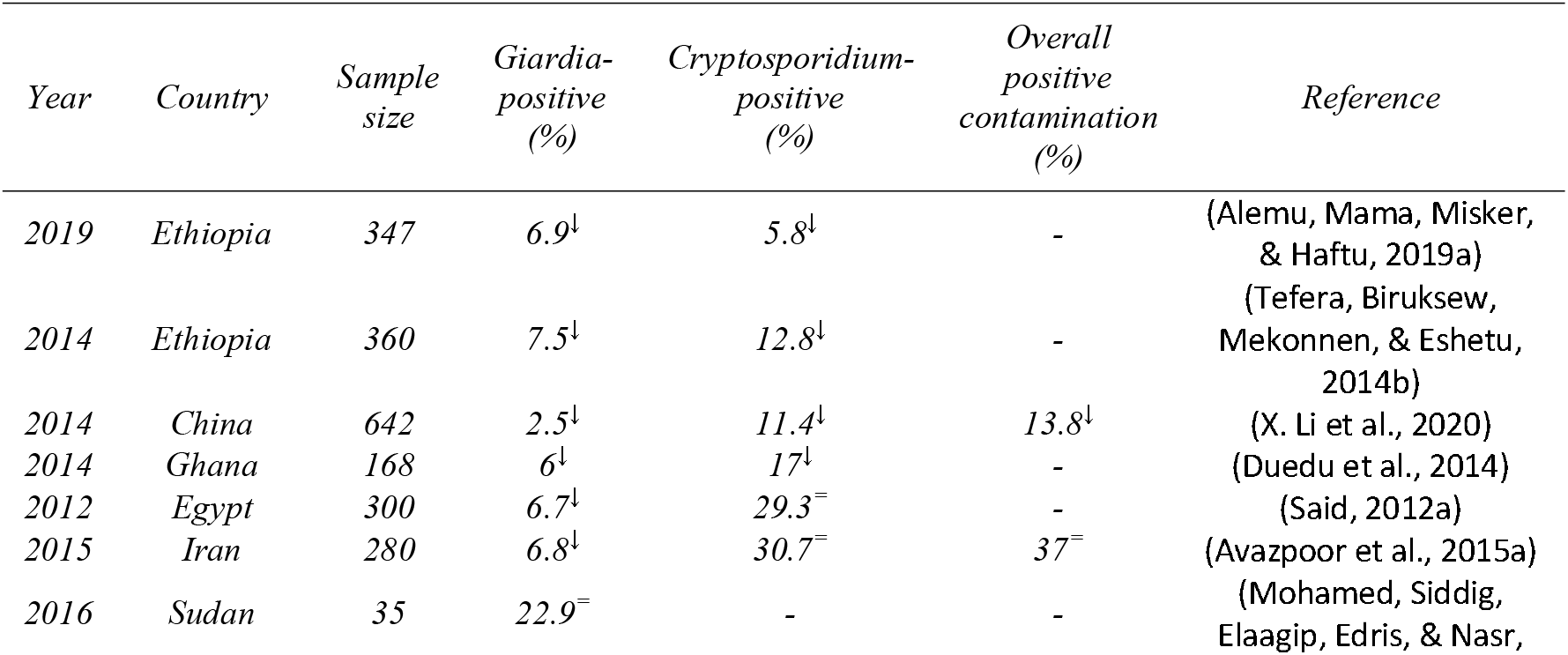
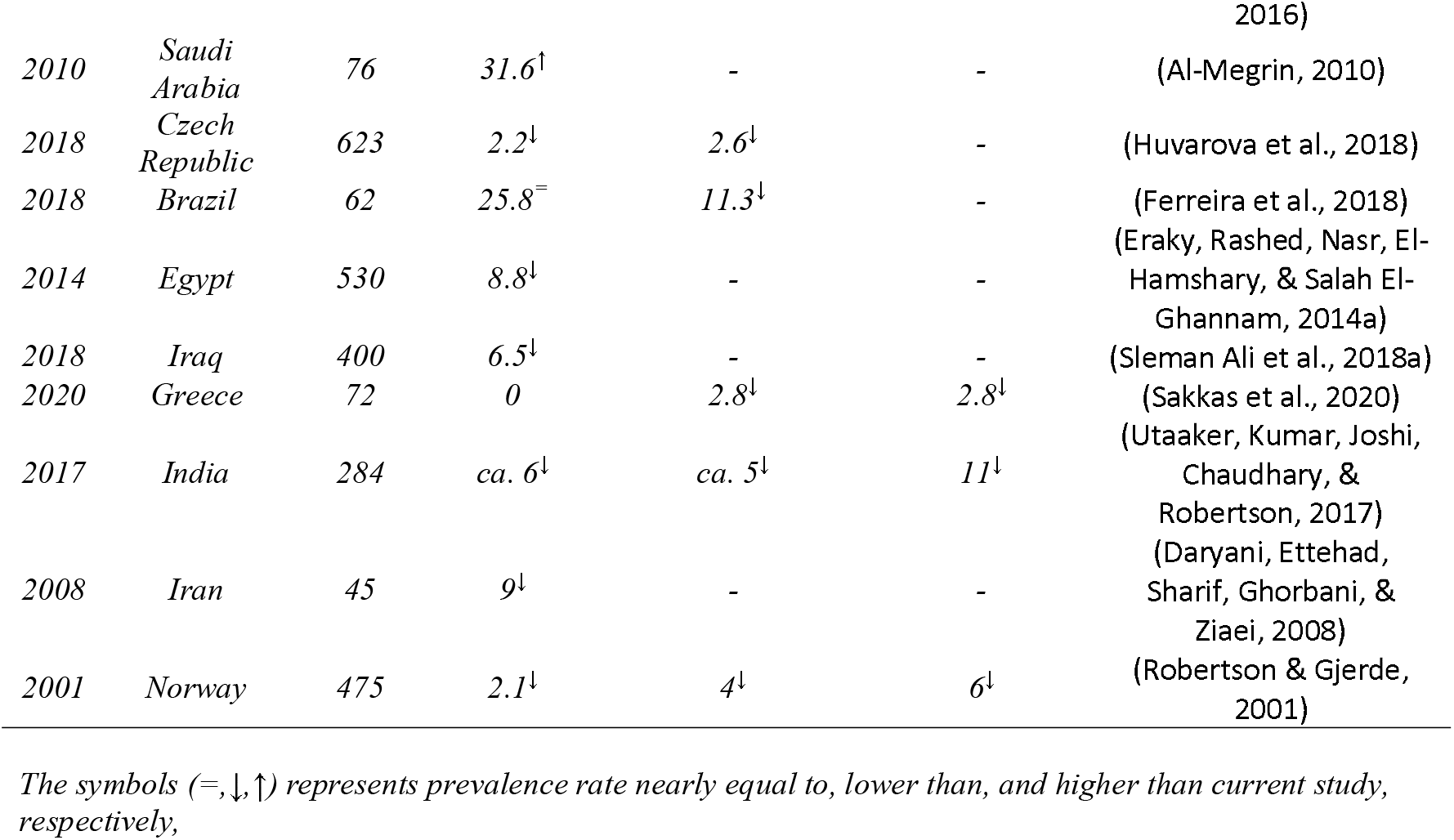
Levels of contamination of vegetable produces by the oo(cysts) reported from various geographical regions.

It is interesting to note that higher number of samples reported in this research were contaminated with *Cryptosporidium* oocysts (33.3%) than *Giardia* (23.2%) (chi-square test: df=1, P<0.05). Similar results were obtained by Robertson and Gjerde (Robertson & Gjerde, 2001) and Moyad Avazpoor (Avazpoor et al., 2015b), in a survey to assess parasitic contamination in various fruits and vegetables. *Cryptosporidium* oocysts adhere strongly to surfaces and may penetrate the stomatal openings of leafy vegetables due to which it is tough to remove them from the surfaces if they are not washed properly (Leitch & He, 2011). It has been found that fertilizing crops with cattle and sheep manure containing viable oocysts of *Cryptosporidium* as oocysts can survive for several months in manure especially at low temperature (Olson, Goh, Phillips, Guselle, & McAllister, 1999; Ryan, Fayer, & Xiao, 2014) may help contaminate the harvest.

*Giardia* cysts have relatively shorter life span in the environment. They can survive at -4 °C in water and in soil for less than a week whereas *Cryptosporidium* oocysts could survive for more than twelve weeks (Olson et al., 1999). In some recorded outbreaks of foodborne transmission of giardiasis, the food handler has been the suspected source in five out of nine cases (Figgatt et al., 2017; Osterholm et al., 1981). Nonetheless, a relatively fewer number in the study suggests that contamination is more probable from water or transporting vehicles than direct contact with an individual.

In our study, 19% vegetable samples were positive for both (oo)cysts (Table S3), which is almost three times (8.3%) the study by Bodhidatta *et al*.(Bodhidatta, 2016). The effect of polyparasitism can aggravate the morbidity rate (Sayasone, Utzinger, Akkhavong, & Odermatt, 2015; Steinmann, Utzinger, Du, & Zhou, 2010). Also, multiple infections can affect individual’s immune system to respond the infection as the synergistic interaction has been observed between co-infecting pathogens and diarrhoeal pathogenesis. A study in Ecuador, South America reported simultaneous infection with rotavirus and *Giardia* that resulted in a greater risk of having diarrhoea than would be expected if the organism acted independently of one another (Bhavnani, Goldstick, Cevallos, Trueba, & Eisenberg, 2012).

### 3.2. Levels of contamination in different types of vegetables

The contamination varied in different vegetable types. Out of 651 vegetable samples, the prevalence rate of at least one cyst ranged from 20% in carrot to 48% in spinach. The samples with only *Giardia* contamination ranged from 5% in carrot to 32% in tomato. Similarly, the prevalence rate of *Cryptosporidium* only ranged from 18% in carrot to 44% in spinach (Figure 2b, Table S2). The prevalence was significantly lower in carrots when compared with spinach, tomatoes, cabbage and green chilli (Pairwise proportional test, P<0.05). Similar results have been reported from other countries (Robertson & Gjerde, 2001; Sleman Ali et al., 2018b).

In another study (Anh, Tram, Klank, Cam, & Dalsgaard, 2007), carrot had a moderate frequency of detection of *Cryptosporidium* parasites as a root crop it is continuously exposed to soil contaminants. We found that tomato had relatively higher contamination ratio than other vegetables tested in contrast to a study reported from Libya (Abougrain, Nahaisi, Madi, Saied, & Ghenghesh, 2010) where only 3% *Giardia* spp. in tomato, 19% in cucumber, and just 4% in lettuce were contaminated. For the root vegetables like carrot, 14% contamination was reported in India (Utaaker et al., 2017), 4% in North Central Nigeria(Amaechi, Ohaeri, Ukpai, & Adegbite, 2016) and 6.4% in Southern Ethiopia(Alemu, Mama, Misker, & Haftu, 2019b). The contamination of leafy greens such as the lettuce was low in Canada (0%)(Lalonde & Gajadhar, 2016) and India (0%) (Utaaker et al., 2017) compared to that reported in Egypt (43.3%)(Said, 2012b).

This result shows that raw vegetables consumed by the Nepali population are often contaminated with intestinal parasites. These types of vegetables should be considered as a potential source of parasitic contamination among general people. These findings underscore the public health implication of consumers of these vegetables being at high risk of infection with giardiasis and cryptosporidiosis. Our study reveals that parasitic contamination rate in vegetables sold from vegetable markets in Nepal is significantly considerable. A significant number of contaminated samples had presence of both (oo)cysts and comparing between the two parasites, *Cryptosporidium* oocysts were more frequently detected than *Giardia* cysts. It was observed that there was significant seasonal variation, whereas contamination rate did not differ in different vegetables and at different locations. The scope of the study did not include viability, infectivity, strain/isolate of parasite and potential routes of contamination such as fertilization procedures, contaminated environments during handling, transport and storage, or direct contamination from individuals involved in the production and processing of products. We recommend the public health sector to advocate consuming vegetables only after adequate washing or proper cooking. In addition, consumers of this region should be informed to properly disinfect these vegetables before consuming them raw.

### 3.3. Distribution of positive samples in different locations

The sampling sites in this research represent Bagmati, Lumbini, Karnali and Sudharpaschim provinces of Nepal. The prevalence rate of vegetable samples at different sites ranged from 13% in Dhading to 61% in Dhangadi. The prevalence of only *Cryptosporidium* oocysts contamination ranged from 7.9% in Dhading to 30.3% in Surkhet. Similarly, the prevalence rate of only *Giardia* cysts contamination ranged from 11.1% in Dhading to 61.1% in Dhangadi (Figure 3A and Table S4).

**Figure 3:**
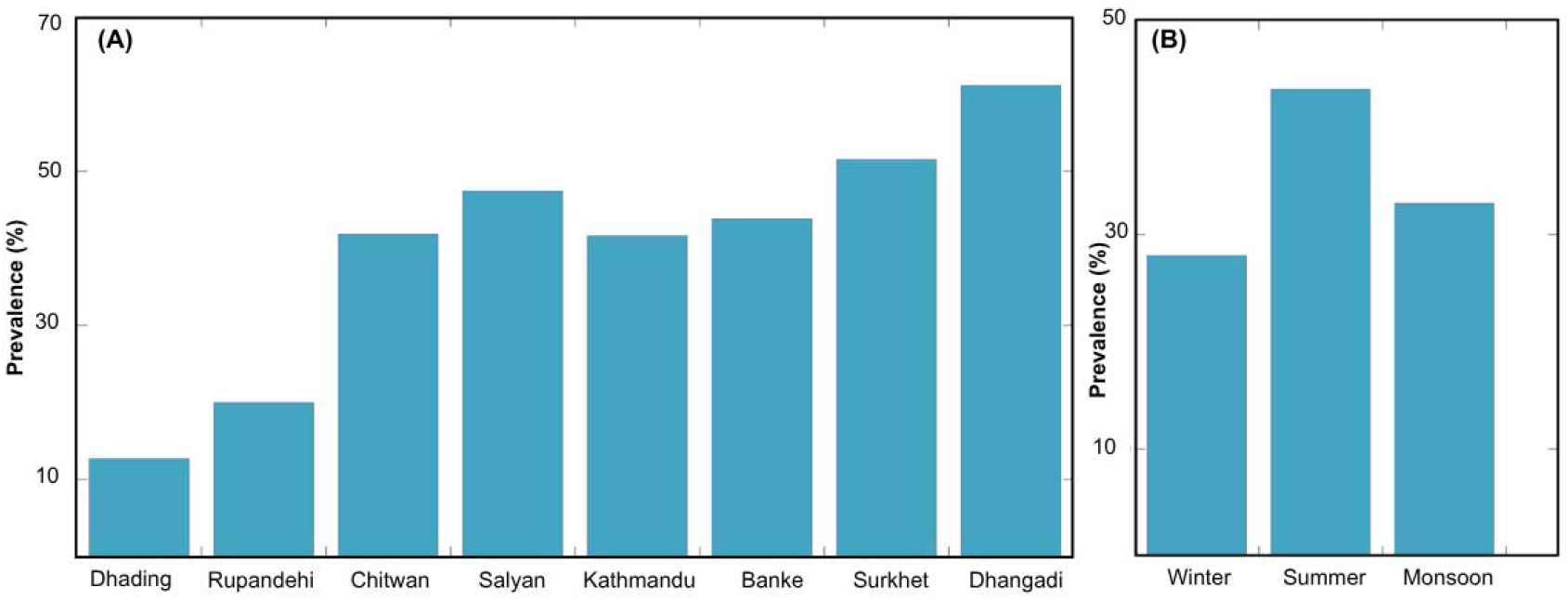
Prevalence of (oo)cysts on vegetable samples collected from different locations (A) and their seasonal variation in the samples collected at Kathmandu (B).

The overall contamination rate (either *Cryptosporidium*, or *Giardia* or both) was not significantly different across different districts except in Dhading where it was significantly lower compared to other locations (Pairwise proportional test, P<0.05). The contamination rate in Rupandehi is lower than in Kathmandu and Dhangadhi. The similar result was observed for the contamination rate of *Cryptosporidium*. In case of *Giardia* cysts, the difference of occurrence in different districts was not significant (Pairwise proportional test, P>0.05). In case of Dhading, most vegetables are directly delivered from local farm to the markets, which minimizes the transportation time and exposure to get contaminated, explaining the lower contamination rate. Also, the water used for washing the vegetables may not be contaminated as the parasitic contamination of produce has been linked to washing by contaminated surface water (J. Li, Wang, Karim, & Zhang, 2020). This result implies that the occurrence of both (oo)cysts did not depend on the locations and there is an equal chance of infection with both (oo)cysts in all locations, which might be due to homogeneous dispersion of parasites and poor sanitary conditions in the community.

### 3.4. Seasonal variation of (oo)cysts contamination

We tested 214 samples from Kalimati fruits and vegetable market, Kathmandu representing summer, monsoon, and winter seasons. Overall, the contamination rate was 28% for winter, 43% for summer and 33% for monsoon seasons. *Cryptosporidium* spp. Only oocysts were detected in 18.7% winter samples, 27.1% monsoon samples, and 36.2% summer samples. Similarly, *Giardia* only cysts were detected in 13.3% samples collected during winter, 27.1% samples during monsoon, and 21.7% samples during summer (*see* Figure 3B and Table S5). In general, the prevalence rate was higher in summer season in Kathmandu, but it was not statistically significant (P>0.05). The prevalence rate of (oo)cysts was significantly lower in winter (Pairwise proportional test, P<0.05) whereas the prevalence of both *Cryptosporidium* oocysts and *Giardia* cysts didn’t vary significantly between summer and monsoon seasons (Pairwise proportional test, P>0.05). Warm and humid condition can aid the survival of (oo)cysts and heavy rainfall, and flooding can promote the transmission of the infections.

A study of seasonal outbreaks of *Cryptosporidium* in different vegetables in Kathmandu, Nepal for three consecutive years from May to October showed a high prevalence of up to 30% during the rainy season (Ghimire, Mishra, & Sherchand, 2005). Warm temperature, humidity and stagnation of open water can increase the exposure to parasites, extend the infective period and transmission of (oo)cysts raising the possibility of contacting with the population (Lal, Baker, Hales, & French, 2013). Frequent use of untreated wastewater for irrigation of vegetable during summer can result in a higher rate of parasitic contamination (Fallah, Makhtumi, & Pirali-Kheirabadi, 2016).

It is usually observed that *Cryptosporidium* oocysts are found in a higher percentage than the cysts in summer and monsoon. According to Amahmid et al., (1999) *Giardia* cysts disappeared after three days because of high temperature, and intense solar radiation. Factors like solar radiation, temperature, humidity, and rainfall directly affect the persistence of microorganisms. A recent review, which analysed information on patterns of important human enteric zoonotic diseases in temperate climatic zones in developed countries, reported that there is a clear peak of cryptosporidiosis cases in summer seasons. In contrast, giardiasis showed a relatively small peak in summer (Abeywardena et al., 2015).

In Nepal, usually a high number of protozoal infections are reported in wet and warm seasons. Faecal disposition especially by the children, near the roadside or along the riverside at the night-time is quite common in rural areas. Cattles freely grazes in these areas. These may contaminate river water and sewage. In the rainy season, the seepage of water from these sources may contaminate vegetables either when they are openly kept in soil for selling or when they are in the fields just before harvesting. Using human waste as manure, irrigating crops with contaminated water, washing the produce with contaminated water could lead to contamination of vegetables with both (oo)cysts. A common source of infection appears to be contaminated water and transmission rates seem to peak between April and September (summer and monsoon) (Ghimire et al., 2005).

## Acknowledgement

We thank Dikshya Puri, Krisha Pokhrel, Sachin Sejuwal, Samiksha Pokhrel, Shishir Pandey and Jhalak Paudel for their help during sample collection and Dr. Ramesh Sapkota for discussion on statistical analysis.

## Supporting Information

Raw data for the number of (oo)cysts observed in all the vegetables collected from nine major vegetable collection sites of Nepal.

