## Supplemental Table 1-9 for "Occurrence of (oo)cysts of *Cryptosporidium* and *Giardia* on vegetables across Nepal"

*Correspondence:

Basant Giri

### Supplementary information

*Table S1: List of samples at different sampling locations*

| Name of samples | Number of samples | | | | | | | | | |
|---|---|---|---|---|---|---|---|---|---|---|
| | Dhading | Butwal | Chitwan | Salyan | Kalimati | Balkhu | Banke | Surkhet | Dhangadi | Total |
| Tomato | 10 | 10 | 10 | 3 | 30 | 30 | 10 | 9 | 5 | 117 |
| Radish | 10 | 10 | 10 | 2 | 29 | 0 | 2 | 4 | 3 | 70 |
| Cabbage | 10 | 10 | 10 | 4 | 30 | 18 | 6 | 7 | 4 | 99 |
| Cucumber | 9 | 10 | 10 | 0 | 30 | 27 | 3 | 5 | 3 | 97 |
| Green onion | 0 | 10 | 0 | 0 | 9 | 0 | 0 | 0 | 0 | 19 |
| Green chilli | 10 | 10 | 10 | 3 | 30 | 30 | 10 | 6 | 3 | 112 |
| Carrot | 10 | 10 | 10 | 3 | 30 | 0 | 1 | 2 | 0 | 66 |
| Spinach | 4 | 10 | 7 | 4 | 26 | 20 | 0 | 0 | 0 | 71 |
| **Total** | **63** | **80** | **67** | **19** | **214** | **125** | **32** | **33** | **18** | **651** |

*Table S2: Prevalence of (oo)cysts in vegetable samples*

| Vegetable name | No. of samples tested | No. of contaminated samples | | |
|---|---|---|---|---|
| | | At least one of (oo)cysts | Giardia only (%) | Cryptosporidium only (%) |
| Tomato | 117 | 53 (45%) | 37 (32%) | 49 (42%) |
| Raddish | 70 | 21(30%) | 12 (17%) | 18 (26%) |
| Cabbage | 99 | 44 (44%) | 26 (26%) | 38 (38%) |
| Cucumber | 97 | 27 (28%) | 18 (19%) | 22 (23%) |
| Green Onion | 19 | 4 (21%) | 2 (11%) | 4 (21%) |
| Green Chilli | 112 | 48 (43%) | 31(28%) | 43 (38%) |
| Carrot | 66 | 13 (20%) | 3 (5%) | 12 (18%) |
| Spinach | 71 | 34 (48%) | 22 (31%) | 31(44%) |
| | Average (%) | 37.5% | 23.2% | 33.3% |

*Table S3: Contamination of vegetables at different sampling locations*

| Vegetables | (oo)cysts types | Sampling districts | | | | | | | | | | | | |
|---|---|---|---|---|---|---|---|---|---|---|---|---|---|---|
| | | Dhading | Rupandehi | Chitwan | Salyan | Kalimati_win | Kalimati_Sum | Kalimati_Mon | Balkhu | Banke | Surkhet | Dhangadi | **Total (count)** | **Total (%)** |
| Tomato | Both | 1 | 1 | 3 | ND | ND | 2 | ND | 17 | 2 | 5 | 2 | **33** | **28.2** |
| | Giardia only | ND | ND | 1 | ND | 1 | ND | 2 | ND | ND | ND | ND | **4** | **3.4** |
| | Crypto only | ND | 1 | 3 | 1 | 2 | 3 | ND | 1 | 1 | 2 | 2 | **16** | **13.7** |
| Radish | Both | ND | 1 | 1 | ND | ND | 1 | 2 | ND | 1 | 2 | 1 | **9** | **12.9** |
| | Giardia only | ND | ND | ND | ND | 2 | ND | 1 | ND | ND | ND | ND | **3** | **4.3** |
| | Crypto only | 2 | ND | 2 | 1 | 1 | 2 | ND | ND | 1 | ND | ND | **9** | **12.9** |
| Cabbage | Both | ND | 1 | ND | 2 | 2 | 1 | 5 | 6 | 1 | 1 | 1 | **20** | **20.2** |
| | Giardia only | ND | ND | ND | ND | 2 | 4 | ND | ND | ND | ND | ND | **6** | **6.1** |
| | Crypto only | 1 | 3 | 2 | 1 | 2 | 1 | 1 | 2 | 2 | 1 | 2 | **18** | **18.2** |
| Cucumber | Both | 1 | ND | ND | ND | 1 | 2 | 2 | 5 | ND | 1 | 1 | **13** | **13.4** |
| | Giardia only | 1 | ND | ND | ND | 1 | 1 | ND | 2 | ND | ND | ND | **5** | **5.2** |
| | Crypto only | ND | 1 | ND | ND | ND | 2 | ND | 3 | 1 | 2 | ND | **9** | **9.3** |
| Green Onion | Both | ND | 1 | ND | ND | ND | 1 | ND | ND | ND | ND | ND | **2** | **10.5** |
| | Giardia only | ND | ND | ND | ND | ND | ND | ND | ND | ND | ND | ND | **0** | **0.0** |
| | Crypto only | ND | 1 | ND | ND | ND | 1 | ND | ND | ND | ND | ND | **2** | **10.5** |
| Green Chilli | Both | 1 | 3 | 4 | 1 | ND | 3 | 2 | 10 | 1 | 1 | ND | **26** | **23.2** |
| | Giardia only | ND | ND | 1 | ND | ND | ND | ND | 1 | 3 | ND | ND | **5** | **4.5** |
| | Crypto only | ND | ND | 3 | 1 | 3 | 3 | 1 | 2 | 1 | 1 | 2 | **17** | **15.2** |
| Carrot | Both | 1 | ND | ND | ND | ND | ND | 1 | ND | ND | ND | ND | **2** | **3.0** |
| | Giardia only | ND | ND | ND | ND | 1 | ND | ND | ND | ND | ND | ND | **1** | **1.5** |
| | Crypto only | ND | ND | 6 | 1 | 1 | 1 | ND | ND | ND | 1 | ND | **10** | **15.2** |
| Spinach | Both | ND | 2 | ND | ND | ND | ND | 3 | 14 | ND | ND | ND | **19** | **26.8** |
| | Giardia only | ND | ND | ND | ND | ND | ND | 1 | 2 | ND | ND | ND | **3** | **4.2** |
| | Crypto only | ND | 1 | 2 | 1 | 2 | 2 | 2 | 2 | ND | ND | ND | **12** | **16.9** |

***Table S4: Distribution of Giardia and Cryptosporidium contamination in different locations of Nepal***

| Sampling location | Number of samples tested | Contaminated samples | | | | | |
|---|---|---|---|---|---|---|---|
| | | At least one (oo)cyst | | Giardia only | | Cryptosporidium only | |
| | | Count | % | Count | % | Count | % |
| Dhading | 63 | 8 | 13% | 5 | 7.9% | 7 | 11.1% |
| Rupandehi | 80 | 16 | 20% | 9 | 11.3% | 16 | 20.0% |
| Chitwan | 67 | 28 | 42% | 10 | 14.9% | 26 | 38.8% |
| Salyan | 19 | 9 | 47% | 3 | 15.8% | 9 | 47.4% |
| Kathmandu | 339 | 141 | 42% | 101 | 29.8% | 120 | 35.4% |
| Banke | 32 | 14 | 44% | 8 | 25.0% | 11 | 34.4% |
| Surkhet | 33 | 17 | 52% | 10 | 30.3% | 17 | 51.5% |
| Dhangadi | 18 | 11 | 61% | 5 | 27.8% | 11 | 61.1% |

***Table S5: Prevalence of (oo)cysts in different seasons in Kathmandu***

| Season | Number of samples tested | Contaminated samples | | | | | |
|---|---|---|---|---|---|---|---|
| | | At least one (oo)cyst | | Giardia only | | Cryptosporidium only | |
| | | Count | % | Count | % | Count | In % |
| Winter | 75 | 21 | 28% | 10 | 13.3% | 14 | 18.7% |
| Summer | 69 | 30 | 43% | 15 | 21.7% | 25 | 36.2% |
| Monsoon | 70 | 23 | 33% | 19 | 27.1% | 19 | 27.1% |

**Statistical analysis**

*Table S6: Chi square significant test for contaminated and non-contaminated sample for different locations, vegetables, and seasons*

| Total contamination | | |
|---|---|---|
| Overall | p-values | Degree of Freedom |
| Location | 2.6E-06 | 7 |
| Vegetable | 4.3E-06 | 7 |
| Season | 1.4E-01 | 2 |

- There are 28 combinations.
- Significant cut off = 0.05, p-cut = 0.05
- After Bonferroni correction significant cut off value = 0.001786

*Table S7: **Pairwise chi square test p-values from different locations (like Dhading-Rupandehi, Dhading-Chitwan, Dhading-Salyan, Dhading-Kathmandu, Dhading-Banke, Dhading-Dhangadi and so on)***

| Sampling locations | Dhading | Rupandehi | Chitwan | Salyan | Kathmandu | Banke | Surkhet | Dhangadi |
|---|---|---|---|---|---|---|---|---|
| Dhading | - | 3.50E-01 | 4.51E-04 | 3.23E-03 | 2.46E-05 | 1.73E-03 | 1.08E-04 | 7.51E-05 |
| Rupandehi | 3.50E-01 | - | 7.09E-03 | 2.97E-02 | 5.39E-04 | 1.99E-02 | 1.79E-03 | 1.21E-03 |
| Chitwan | 4.51E-04 | 7.09E-03 | - | 8.64E-01 | 1.00E+00 | 1.00E+00 | 4.81E-01 | 2.32E-01 |
| Salyan | 3.23E-03 | 2.97E-02 | 8.64E-01 | - | 7.97E-01 | 1.00E+00 | 1.00E+00 | 6.11E-01 |
| Kathmandu | 2.46E-05 | 5.39E-04 | 1.00E+00 | 7.97E-01 | - | 9.61E-01 | 3.59E-01 | 1.65E-01 |
| Banke | 1.73E-03 | 1.99E-02 | 1.00E+00 | 1.00E+00 | 9.61E-01 | - | 7.05E-01 | 3.77E-01 |
| Surkhet | 1.08E-04 | 1.79E-03 | 4.81E-01 | 1.00E+00 | 3.59E-01 | 7.05E-01 | - | 7.16E-01 |
| Dhangadi | 7.51E-05 | 1.21E-03 | 2.32E-01 | 6.11E-01 | 1.65E-01 | 3.77E-01 | 7.16E-01 | - |

* There are 28 combinations

*The significant p-value cut off is 0.05. After Bonferroni correction it is adjusted to 0.0018

*Table S8: Pairwise chi square test p-values from different vegetables (Tomato-Radish, Tomato-Cabbage, Tomato-Cucumber, Tomato-Green Onion, Tomato-Green Chilli, Tomato-Carrot, Tomato-Spinach and so on*

| | Tomato | Radish | Cabbage | Cucumber | Green Onion | Green Chilli | Carrot | Spinach |
|---|---|---|---|---|---|---|---|---|
| Tomato | - | 0.0554 | 1.0000 | 0.0129 | 0.0825 | 0.8112 | 0.0010 | 0.8460 |
| Radish | 0.0554 | - | 0.0817 | 0.8952 | 0.6299 | 0.1136 | 0.2346 | 0.0450 |
| Cabbage | 1.0000 | 0.0817 | - | 0.0232 | 0.0997 | 0.9259 | 0.0019 | 0.7732 |
| Cucumber | 0.0129 | 0.8952 | 0.0232 | - | 0.7433 | 0.0346 | 0.3174 | 0.0122 |
| Green Onion | 0.0825 | 0.6299 | 0.0997 | 0.7433 | - | 0.1229 | 1.0000 | 0.0655 |
| Green Chilli | 0.8112 | 0.1136 | 0.9259 | 0.0346 | 0.1229 | - | 0.0029 | 0.6071 |
| Carrot | 0.0010 | 0.2346 | 0.0019 | 0.3174 | 1.0000 | 0.0029 | - | 0.0010 |
| Spinach | 0.8460 | 0.0450 | 0.7732 | 0.0122 | 0.0655 | 0.6071 | 0.0010 | - |

*There are 28 combinations
*The significant p-value cut off is 0.05, considering Bonferroni correction it is adjusted to 0.0018

*Table S9: Pairwise chi square test p-values from different seasons*

| | Winter | Summer | Monsoon |
|---|---|---|---|
| Winter | - | 0.0524 | 0.5250 |
| Summer | 0.0524 | - | 0.1974 |
| Monsoon | 0.5250 | 0.1974 | - |

*There are 3 combinations
*The significant p-value cut off is 0.05, considering Bonferroni correction it is adjusted to 0.0167
